# Phylogenetic inference from an incomplete fossil record

**DOI:** 10.64898/2026.06.24.734220

**Authors:** Niklas Hohmann, Rachel C. M. Warnock, Emilia Jarochowska

## Abstract

Fossil data is crucial to construct phylogenetic time trees, which serve as the basis to test a wide range of evolutionary hypotheses. While the fossil record is known to be incomplete, modern stratigraphy provides predictions of the structure of the fossil record as expressed by gap location and duration. Advances in phylogenetic model development allow us to propagate this information into Bayesian phylogenetic inference in the form of priors on time-variable fossil sampling. However, the impact and role of stratigraphic architectures on time tree inference has so far remained unexplored.

We introduce a novel simulation framework that combines realistic stratigraphic forward models with phylogenetic simulations. Using this framework, we examine (1) how stratigraphically plausible model violations of fossil sampling due to gaps affect total-evidence inference under the fossilized birth-death model and (2) if stratigraphic knowledge on gap duration and timing improves inference when incorporated in priors on fossil sampling.

We find that total-evidence analysis is robust to stratigraphically plausible distribution of gaps in disparate stratigraphic architectures, with results being instead dominated by the number of morphological characters. Surprisingly, incorporating information on prominent gaps in the stratigraphic record does not improve phylogenetic inference.

Our results suggest that phylogenetic inference is robust to model violations introduced by stratigraphic gaps over short timescales, with results being dominated by a priori known data availability constraints such as morphological character matrix size. This research establishes the foundations for joint modeling of phylogenetic and stratigraphic processes and narrows the knowledge gap between paleontology, stratigraphy, and neontology.

## Introduction

Morphological and temporal data from fossils provide direct physical evidence for past evolutionary change beyond timescales accessible to direct human observation. While there remains some skepticism towards the utility of fossil data due to the incompleteness of the fossil record, paleontologists usually take a positivist stance, arguing that the fossil record is suitable to test a wide range of evolutionary hypotheses (Darwin 1859; Paul 1992; Springer 1995; Kidwell and Holland 2002; Benton and Emerson 2007; Escapa and Pol 2011).

Fossils are embedded in sedimentary rocks, and removal of previously deposited sediment or nondeposition of new sediment results in gaps in both the rock and, by extension, the fossil record. While the resulting stratigraphic incompleteness (proportion of time preserved in the rock record) was long considered problematic, progress in stratigraphic forward modeling has demonstrated that the *relative* contributions of short vs. long gaps determines how well evolutionary history can be recovered (Hohmann et al. 2024). Spatial and temporal extent of gaps and their frequency are determined by external forcing on stratigraphic architectures (Holland and Patzkowsky 1999). On macroevolutionary timescales, global controls on eustasy and tectonics can lead to the systematic removal of fossils from particular intervals of the Earths’ history and differences in the fossil record between sedimentary basins.

Historically, the incorporation of stratigraphic and temporal data into phylogenetic inference has been controversial. For example, Smith (2000) argued that “stratigraphic data needlessly inflates the potential sources of error associated with phylogenetic reconstruction”, sparking strong opposition from the paleontological community (Alroy 2002; Fisher et al. 2002) as part of the stratocladistics debate (Wagner 1995; Donoghue 2001; Dzik 2005; Fisher 2008). With the advent of model-based phylogenetic inference, calls have been made to incorporate stratigraphic information more formally into phylogenetic models (Holland 2016; Holland et al. 2025), but only a few examples are available (Wagner and Marcot 2013; Sansom et al. 2015).

Only the introduction of the fossilized birth-death (FBD) model family (Stadler 2010; Didier et al. 2012; Heath et al. 2014; Stadler et al. 2018) made it possible to integrate fossils in a statistically coherent way into phylogenetic models via a constant or piecewise constant fossil sampling process. The FBD model has been applied with great success to empirical datasets and has been thoroughly tested in simulation studies (Wright et al. 2022; Zhang et al. 2023; Mulvey et al. 2025). While the assumption of constant or piecewise constant fossil sampling is guaranteed to be violated in empirical datasets (Holland 2016), the FBD model allows us to directly incorporate stratigraphic information on location and duration of gaps in the form of priors on fossil sampling into Bayesian phylogenetic inference.

Here, we present the first integration of stratigraphic forward models with phylogenetic simulations to explore how the structure of the fossil record influences phylogenetic inference. As an example of the fossil record, our study investigates fossil data preserved in carbonate platforms – environments widely sampled for fossils and recognized as diversity hotspots. We combine models of carbonate platform growth with total-evidence analysis of synthetic fossil records to examine whether:

1. Stratigraphically realistic violations of model assumptions on fossil sampling due to gaps in the fossil record biases total-evidence analysis under the FBD model
2. Incorporation of information on major gaps in the fossil record as priors on fossil sampling improves phylogenetic inference.

## Results

By comparing total evidence inference across differently structured fossil records, we found that the presence of many short gaps (MSG) or a few long gaps (FLG) did not diminish inference quality relative to a continuously sampled fossil record when the total number of fossils is kept constant. Incorporating information on long gaps (> 20% of the observed time interval) as priors into the inference did not improve recovery of model parameters, tree topology, or divergence time estimates.

### Simulated stratigraphic architectures

Simulations of carbonate platforms under both sea level curves (high-amplitude, high frequency and low amplitude, low frequency) yield empirically realistic geometries over the simulated 5 Myr (Williams et al. 2011) (Figure 1). Both under the high frequency, high amplitude sea level curve (scenario 1, Miller et al. (2020)) and the low frequency, low amplitude sea level curve (scenario 2), the simulated carbonate platforms prograde rapidly from their initial topography over the course of the simulation (Figure 1). In scenario 1, a ramp-like geometry develops with many short coeval gaps due to frequent flooding of the platform top (Figure 1 first column, Supplementary Figure 1). At the sampling location 3 km from shore, stratigraphic completeness (percentage of time preserved) is 41% when measured on the timescale of 1 kyr (model resolution), with the majority of gap durations being of the same order of magnitude as the period of the sea level changes (median gap duration 22 kyr, interquartile range (IQR) 32 kyr). No gaps are longer than 300 kyr (Figure 2, Supplementary Table 2), resulting in a stratigraphic record characterized by many short gaps (MSG). In scenario 2, a platform geometry with pronounced separation into platform top and slope deposits develops after an initial phase of progradation, with long gaps on the platform top due to subaerial exposure during drops in sea level (Figure 1 second column, Supplementary Figure 2). At the sampling location, stratigraphic completeness is 27%, with a median gap duration of 49 kyr (IQR: 52 kyr). Two long (> 1 Myr) gaps from 0.425 to 1.606 Myr and 2.628 to 4.191 Myr are present, resulting in a fossil record with few long gaps (FLG) (Figure 2, Supplementary Table 2).

**Figure 1:**
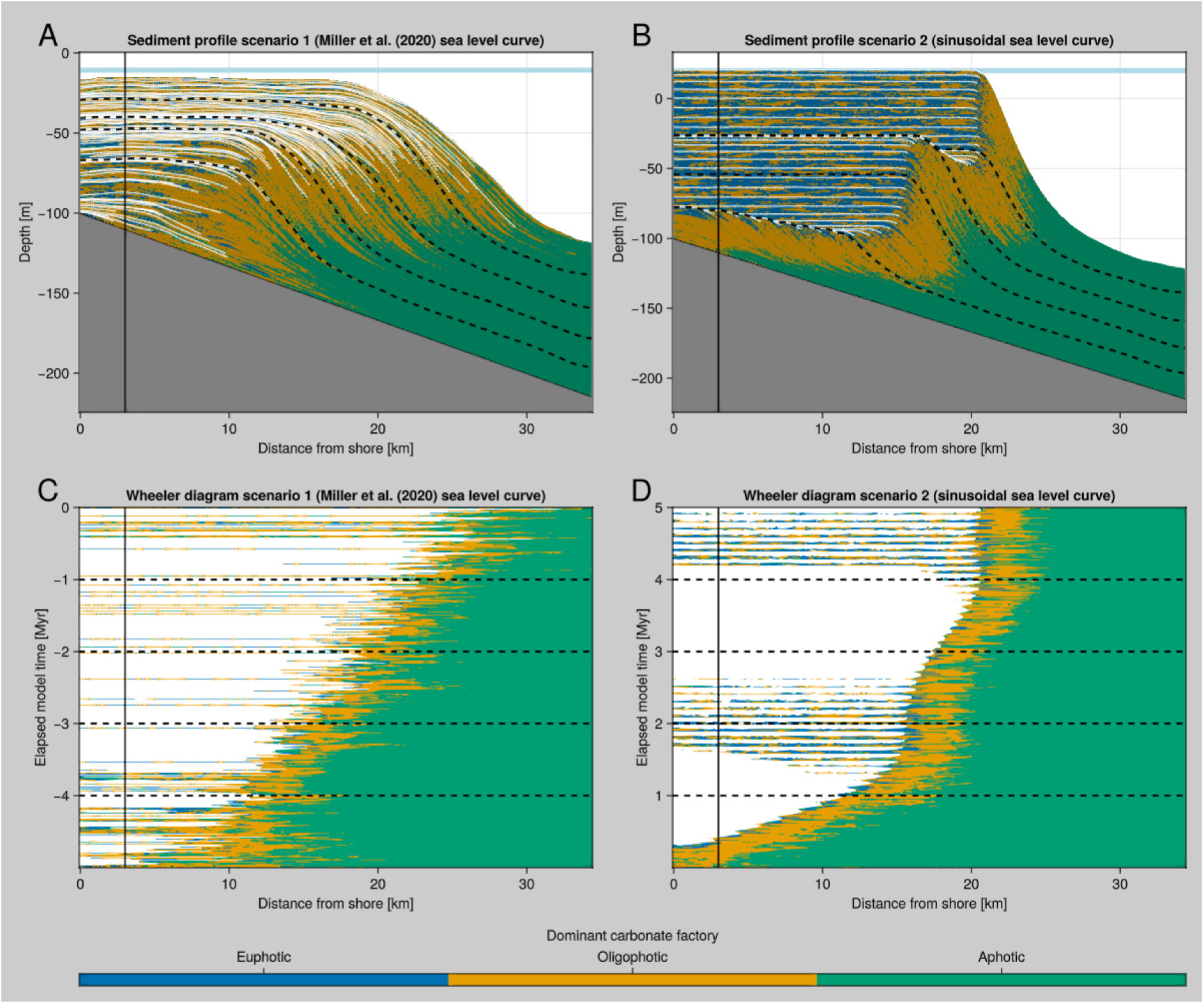
Transects (first row, A and B) and Wheeler (chronostratigraphic) diagrams (second row, C and D) of the attached carbonate platforms simulated under the Miller et al. (2020) sea level (scenario 1, first column, A and C) and the sinusoidal sea level curve (scenario 2, second column, B and D). White lines in the transect are unconformities; white areas in the Wheeler diagram are gaps in the stratigraphic record due to nondeposition and erosion. The solid line indicates the sampling location 3 km offshore at which the fossil records were simulated (Figure 2), dashed lines are coeval lines separating deposits formed over 1 Myr.

**Figure 2:**
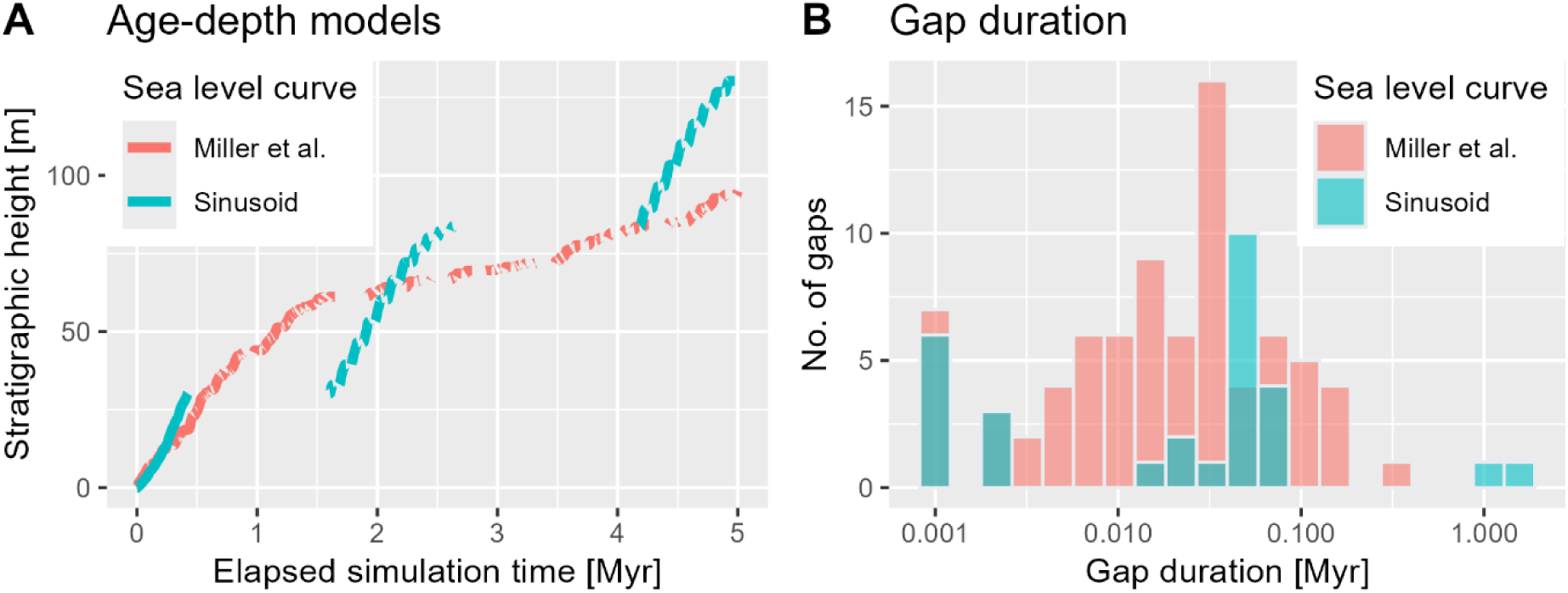
Age-depth models (A) and distribution of gap durations (B) 3 km from shore in the simulated carbonate platforms. See Supplementary Table 2 for gap statistics.

### Inference under stratigraphic model violations

Combining stratigraphically agnostic FBD inference with synthetic fossil records of different structures, we found that stratigraphically plausible model violations of fossil sampling did not degrade results from total-evidence analysis. When running inference under a constant rate FBD (cFBD) model, results did not differ substantially whether synthetic fossil records have continuous fossil sampling (CFS), many short gaps (MSG, scenario 1) or few, but long gaps (FLG, scenario 2) when keeping the number of sampled fossils fixed. This holds for macroevolutionary model parameters (origination and extinction rates, Figure 3 A and B), morphological and molecular clock rates (Figure 3 C and D), origin time (Figure 3 E), tree topology (Figure 4 A), divergence time precision and coverage (Figure 4 B and C) and identification of sampled ancestors (Figure 4 D). The number of available morphological characters was the major determinant of tree topology and divergence time precision and coverage, as well as identification frequency of sampled ancestors (Figure 4).

**Figure 3:**
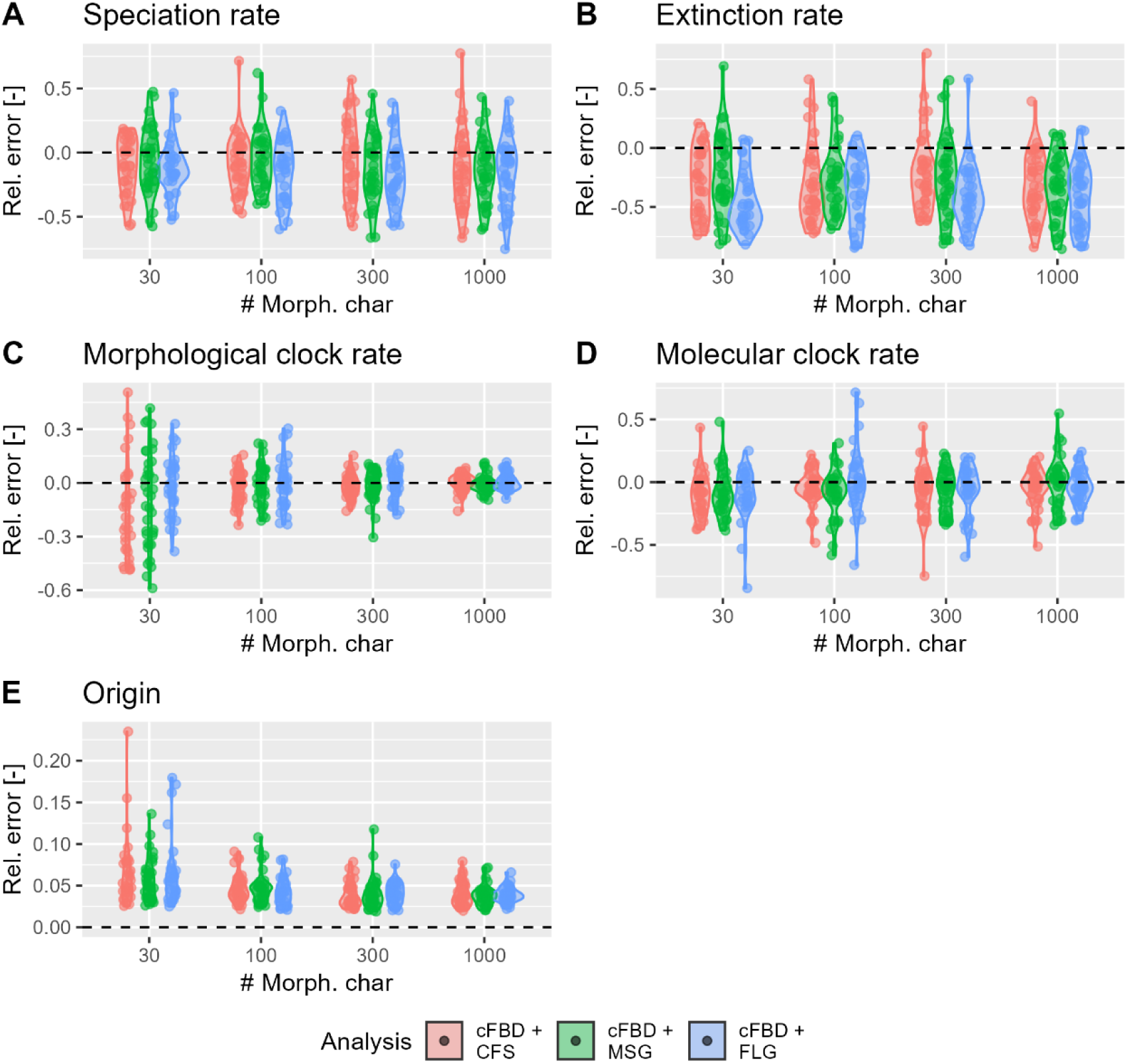
Recovery of model parameters from synthetic fossil records with different structures. Inference is run under the constant rate FBD model (cFBD), applied to a fossil record with constant fossil sampling (CFS), many short gaps (MSG, scenario 1) or few, but long gaps (FLG, scenario 2).

**Figure 4:**
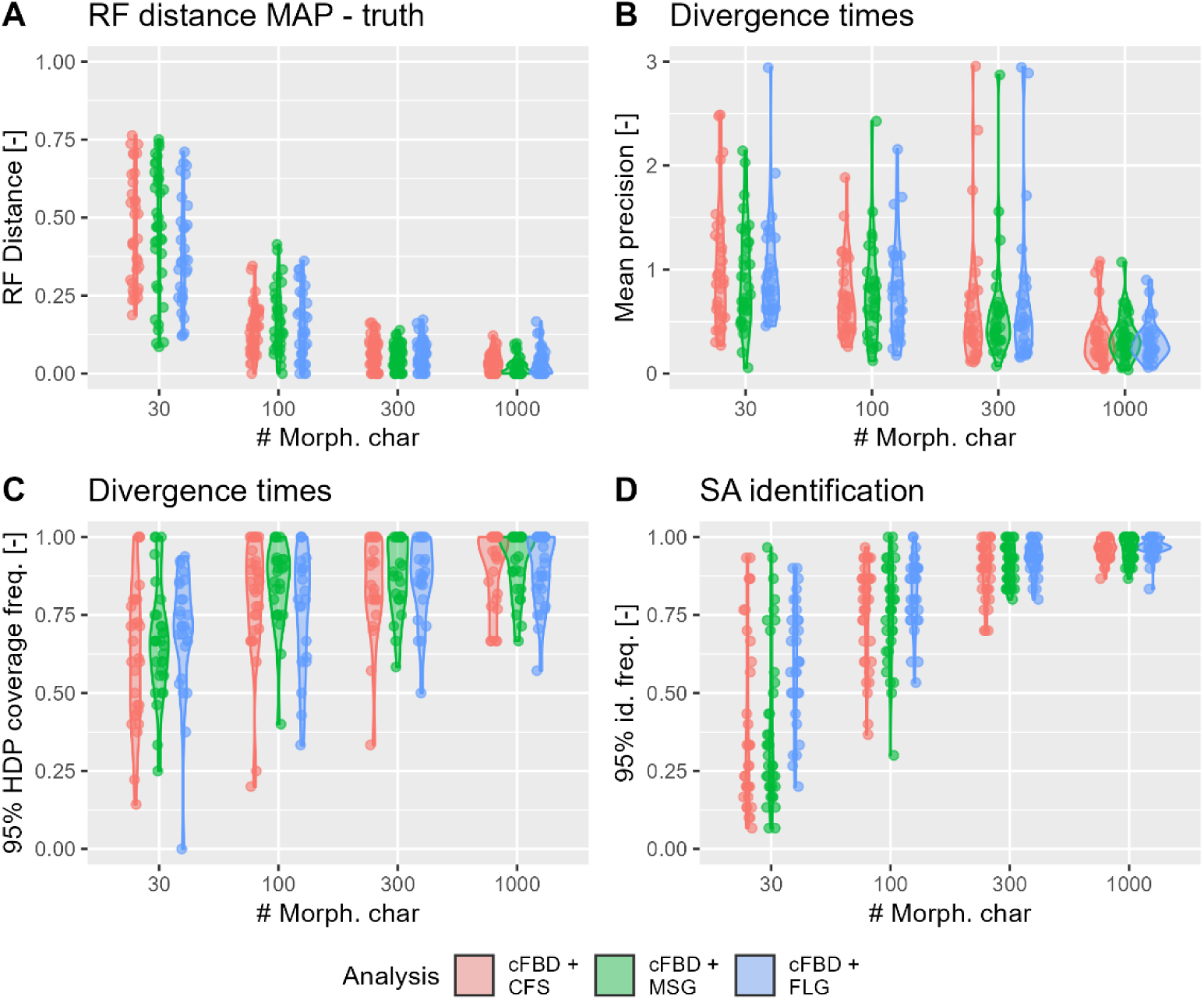
Recovery of (time) tree properties from synthetic fossil records with different structures. Inference is run under the constant rate FBD model (cFBD), applied to a fossil record with constant fossil sampling (CFS), many short gaps (MSG, scenario 1) or few, but long gaps (FLG, scenario 2).

For small to intermediate-sized morphological character matrices (30 to 100 characters), model violations on fossil sampling introduced by gaps in the record worsened the MCMC convergence, with the convergence frequency dropping from 95 to 77.5% between cFBD + CFS and cFBD + FLG (Supplementary Figure 3). For converged runs, coverage and precision for the origination rate was good throughout (Supplementary Figures 4 and 5), while coverage for extinction rate dropped with increasing model violations on fossil sampling due to increases in precision (Supplementary Figure 4 A and B, Supplementary Figure 5). However, the relationship between coverage and fossil record structure was not consistent, e.g., coverage of the morphological clock parameter for small character matrices was better in the presence of long gaps (Supplementary Figure 4 D, Supplementary Figure 7).

**Figure 5:**
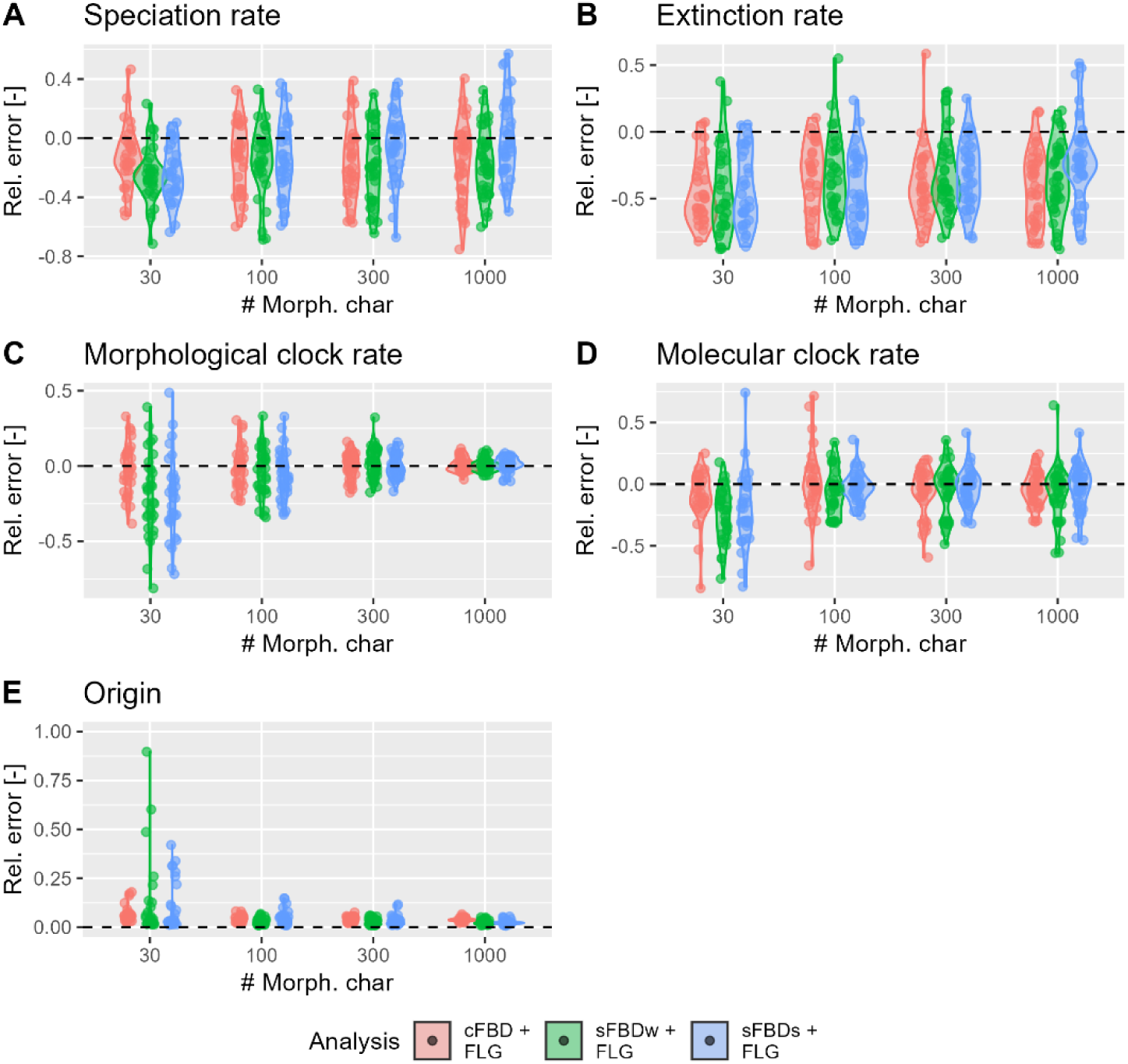
Recovery of model parameters from a synthetic fossil record with few, long gaps (FLG, scenario 2) when incorporating different degrees of stratigraphic knowledge. Inference was run under the stratigraphically agnostic constant rate FBD model (cFBD), weak stratigraphic knowledge using the skyline FBD model with stratigraphically informed breakpoints on fossil sampling (sFBDw), and strong stratigraphic knowledge using the skyline FBD model fixing fossil sampling during gaps to a small numeric value (sFBDs).

### Incorporating stratigraphic information in total evidence analysis

Using a synthetic fossil record with a stratigraphic completeness of 27%, we found that incorporating stratigraphic knowledge about long (> 20% of simulation time) gaps in the form of priors on fossil sampling using the FBD skyline model did not improve estimates from total-evidence analysis. Neither phylogenetic inference with weak stratigraphic information (stratigraphy-informed breakpoints in fossil sampling, sFBDw inference) nor strong stratigraphic information (priors on fossil sampling fixed to a small numeric value during known long gaps, sFBDs inference) performs better than a stratigraphically agnostic inference (constant rate FBD inference, cFBD) when applied to a fossil record with few long gaps (FLG, scenario 2) (Figure 5, Figure 6). This holds for macroevolutionary parameters (origination and extinction rates, Figure 5

**Figure 6:**
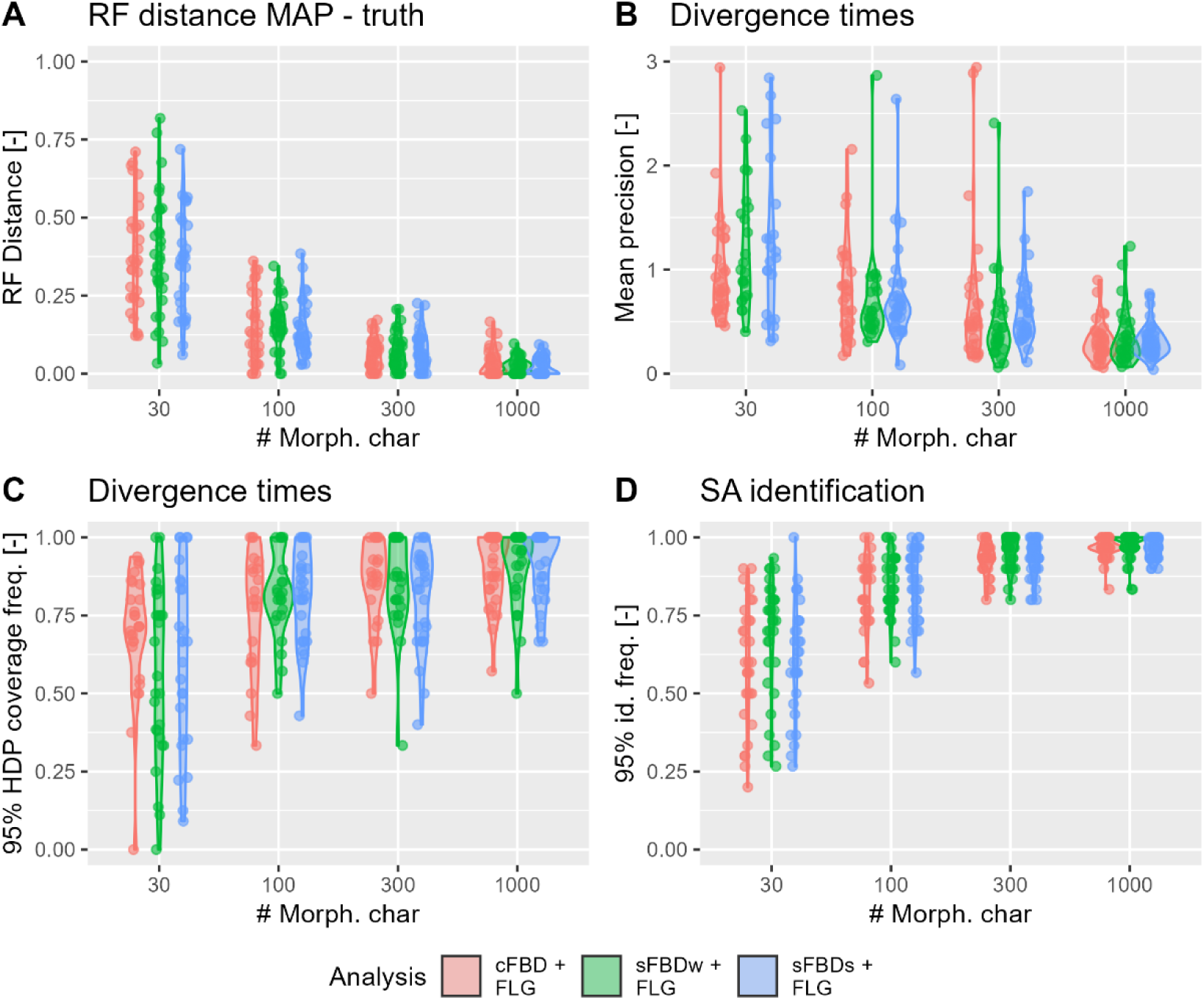
Recovery of (time) tree properties from a synthetic fossil record with few, long gaps (FLG, scenario 2) when incorporating different degrees of stratigraphic knowledge. Inference was run under the stratigraphically agnostic constant rate FBD model (cFBD), weak stratigraphic knowledge using the skyline FBD model with stratigraphically informed breakpoints on fossil sampling (sFBDw), and strong stratigraphic knowledge using the skyline FBD model with fixing fossil sampling during gaps to a small numeric value (sFBDs).

A and B), clock rates (Figure 5, C and D), origin time (Figure 5 E), tree topology (Figure 6 A), divergence time precision and coverage (Figure 6 B and C), and identification of sampled ancestors (Figure 6 D). The exception is the recovery of the origin time for 30 morphological characters, where adding stratigraphic information inflates relative error. Extinction rate is underestimated throughout all analysis variants. The number of available morphological characters is the major determinant of tree topology, divergence time precision and coverage, as well as identification frequency of sampled ancestors (Figure 6).

For character matrices of intermediate size, convergence improves with the incorporation of stratigraphic knowledge, with the biggest gains for character matrices of size 100 (increase in convergence frequency from 77.5 to 87.5 % from cFBD + FLG to sFBDs + FLG, Supplementary Figure 3). Coverage of model parameters is dominated by morphological character matrix size rather than inference setup, although incorporation of stratigraphic knowledge improves the coverage of extinction rate for moderate to large character matrices due to reduced precision (Supplementary Figure 4, Supplementary Figure 6).

## Discussion

Gaps due to erosion and nondeposition of sediment are ubiquitous features of the sedimentary rock record, and, by extension, the fossil record (Kabanov 2017; Tipper 2015). While gap location and duration are difficult to constrain empirically (Tomašových et al. 2022; Davies et al. 2019), they follow predictable patterns within a sequence stratigraphic framework as a function of external drivers such as sea level and tectonics (Catuneanu 2022). This results in an incomplete, but highly structured fossil record (Holland 2017). Our results showed that the structure of the fossil record generated under our simulation scenarios do not affect results of total-evidence analysis under the FBD model. Estimates of macroevolutionary parameters, tree topology, divergence times and origin time were not affected by model violations due to heterogeneous fossil sampling, irrespective of the presence of many short gaps (scenario 1, stratigraphic completeness 41%) or few, long (> 20% simulation time) gaps (scenario 2, stratigraphic completeness 27%) (Figure 3, Figure 4). This suggests that the constant rate FBD model can absorb stratigraphically plausible model violations of fossil sampling in total evidence analyses under the range of simulation scenarios explored here. This result is in line with findings by Zhang et al. (2023) who found that the FBD model is robust to violations of fossil sampling in total-evidence analyses under some scenarios.

Stratigraphic paleobiology posits that the fossil record is a joint expression of biotic and stratigraphic change, and stratigraphic context must be considered when inferring biological change from the fossil record (Holland et al. 2025). Surprisingly, we found that incorporating stratigraphic information in the form of gap position and duration as sampling priors into total evidence analysis did not improve the accuracy of parameter estimates (Figure 5, Figure 6). We attribute this to the robustness of the cFBD model to model violations of fossil sampling, as reduction of model violations (inference under sFBDs instead of cFBD) did not improve performance. Our results indicate that, while reducing model violations by implementing time-dependent sampling rates does not necessarily improve inference results, it can potentially improve convergence (Supplementary Figure 3), even though model complexity is increased (Barido-Sottani et al. 2024).

Throughout the literature, the “incompleteness of the fossil record” serves as an umbrella term for a wide range of stratigraphic, ecological, taphonomic, and socioeconomical effects that differ in how readily they can be identified, analyzed, and accounted for. This includes, but is not limited to, the number of fossil specimens available for analysis, size of morphological character matrices, proportion of time preserved in the fossil record, and spatio-temporal sampling biases due to difference in outcrop area or sampling effort (Donoghue 2001; Woolley et al. 2025; Capobianco 2025; Flannery Sutherland et al. 2019). This study focused on the structure of the fossil record in a narrow sense (duration and location of gaps) rather than established measures of incompleteness (Holland 2017). In our simulations, we fixed the number of morphological characters and fossil to isolate the effect of gaps. Running inference from synthetic data simulated under known data availability constraints makes the effects of character matrix size and number of fossils easily analytically traceable. The resulting “best case” scenario for statistical power and accuracy and precision of the results provides a baseline against which empirical datasets can be compared. In contrast, identifying gaps in the stratigraphic record and constraining their duration is extremely challenging and associated with large uncertainties (Davies et al. 2019), making them effectively epistemic uncertainty in most phylogenetic contexts. For both questions addressed, our results are dominated by the size of the morphological character matrix rather than the structure of the fossil record, stratigraphic incompleteness, or degree of stratigraphic knowledge incorporated. This suggests that epistemic uncertainty originating from incomplete knowledge of stratigraphic incompleteness or the structure of the fossil record plays a subordinate role in phylogenetic inference compared to analytically quantifiable uncertainty rooted in data availability or fossil age uncertainty (Barido-Sottani, Aguirre-Fernández, et al. 2019; Capobianco 2025).

Fossil ages are generally associated with uncertainty, and failure to propagate this uncertainty into FBD analyses is known to result in inaccurate estimates of tree topology and divergence times (Barido-Sottani, Aguirre-Fernández, et al. 2019; Barido-Sottani et al. 2020). To isolate the role of stratigraphic effects and reduce confounding factors, we did not incorporate age uncertainty into our modeling and inference setup. While this remains untested, our expectation is that the effects of age uncertainty are compounding and should always be incorporated alongside other stratigraphic knowledge. Stratigraphic modeling predicts that, in addition to age uncertainty (lack of age precision), fossil ages are systematically biased (not accurate) due to imperfect knowledge of the underlying age-depth relationships (Hohmann, Bickerton, et al. 2026). Not recognizing the unconformity and condensation bias responsible for this effect (sensu (Holland 2000)) would lead to a simultaneous elevation of all observed rates, effectively confounding biological signal and stratigraphic artefacts (Holland and Patzkowsky 2015; Hohmann 2021). Specifically, using the episodic FBD process (Magee and Höhna 2021) will lead to the recovery of elevated origination and extinction rates across major unconformities when fossil ages are derived from simplified age-depth relationships. The extent of this effect will strongly depend on the spatial and temporal scale of the study and its associated depositional system, as well as the availability of absolute age constraints. Especially in deep-time, where radioisotope age constraints are rare and ages commonly are based on biozonation and biostratigraphic correlation, this might produce a dynamic interaction between the structure of stratigraphic architectures and available age constraints and uncertainty.

Our simulation and inference setup was intentionally kept simple to isolate effects resulting from gaps from those of due to other aspects of model complexity or violation (e.g., morphological character encoding, Khakurel et al. (2024)) and to reduce the number of confounding factors. As the most common application of the FBD model that incorporates fossil morphology, we have focused on total evidence analysis, where the molecular partition contributes a lot of information to tree topology (Ronquist et al. 2012; Zhang et al. 2016; Mulvey et al. 2025). It remains an open question how gaps in the fossil record affect inference under more complex simulation setups, fossil-only inference of fully extinct trees (Turner et al. 2017) or how they interact with more complex members of the FBD model family, such as the multitype FBD model (Barido-Sottani and Morlon 2026).

With an interval of 5 Myr covered, this study is at the lower end of the macroevolutionary timescales to which phylogenetics is usually applied. Growth of carbonate platforms over this timescale is well understood and the available simulation tools are mature (Hidding et al. 2025). In addition, focusing on this timescale facilitates comparison of the ways stratigraphy records and modifies different types of palaeobiological information (e.g., mass extinctions and phenotypic evolution (Holland 2000; Hannisdal 2006; Loughney and Holland 2026)) at the level of individual sections. Forward models along onshore-offshore transects on this timescale have been applied with great success to the study of mass extinctions and their recovery, with empirical studies confirming predictions made *in silico* (Danise and Holland 2017; Nawrot et al. 2018; Zimmt et al. 2021; Holland et al. 2025). However, extending the concepts of sequence stratigraphy underlying this body of work to macroevolutionary timescales and larger spatial scales requires more effort than simply running forward models for longer time intervals. Phylogenetic studies typically combine fossil evidence from different sedimentary basins and over timescales potentially larger than the lifespan of any individual basin involved. As a result, there is no single external forcing that controls the structure of the fossil record on this scale (as sea level does in this study), but rather a combination of forcing mechanisms acting on different spatial and temporal scales (Holland 2016), compounded by autigenic dynamics (Liu et al. 2026). Combining evidence across spatial scales involves stratigraphic correlation between sampling sites to place fossils on a joint time scale, a process that is not systematically codified and will introduce additional uncertainties. In addition, gaps are spatially heterogeneous (i.e., time intervals might be preserved in some locations, but not others (Straub et al. 2020)). How the spatiotemporal structure of the fossil record translates into gaps and sampling rates on a joint time scale is an open transdiciplinary challenge that bridges paleobiological and stratigraphic methodologies.

## Methods

### Stratigraphic simulation

We simulated the development of an attached carbonate platform over 5 Myr under two different sea level scenarios in Julia using CarboKitten.jl (Bezanson et al. 2017; Hidding et al. 2025) (Figure 1). Other than the sea level curve, all model parameters were kept identical between scenarios (see Supplementary Information, Supplementary Table 1). In scenario 1, the last 5 Myr of the sea level curve provided by Miller et al. (2020) with orbitally paced high frequency and high amplitude oscillations reflective of icehouse conditions was used (Supplementary Figure 1). In scenario 2, a sinusoidal sea level curve comprised of 3^rd^ and 5^th^ order fluctuations with periods of 2.5 Myr and 0.1 Myr and periods of 20 m and 2 m, respectively, was used (Supplementary Figure 2). Up to the subsidence of 20 m/Myr, this setup is similar to the groundbreaking study on siliciclastic systems by Holland and Patzkowsky (1999). Simulations were run in a 4.5 km (strike, parallel to shore) by 34.5 km (dip, perpendicular to shore) box subdivided into cells of 150 m with time steps of 100 years, recording data every 1000 years (model resolution). From both scenarios, age-depth models, as well as location and timing of gaps from a slice through the middle of the simulation box (along dip) were extracted at 3 km from shore using the R package admtools in R version 4.5.2 (Hohmann et al. 2025; R Core Team 2025) (Figure 2). Stratigraphic completeness (proportion of time preserved) measured at model resolution and gap statistics (quantiles) were calculated from the distribution of gap durations (Supplementary Table 2). Long gaps (longer than 20% of the simulation time, i.e., 1 Myr) were only present under the sinusoidal sea level (scenario 2). These gaps were extracted from the age-depth models, converted from elapsed model time (Myr) into time before present (Ma), and then provided as input to the FBD skyline analysis as fossil sampling rate breakpoints for the sFBDw and sFBDs analysis (see below).

### Phylogenetic simulation

Fossilized birth-death (FBD) trees were simulated conditioned on the duration of the simulation (5 Myr) via the TreeSim package (Stadler 2010), rejecting trees that went extinct before the present. We used an extinction rate of 0.6 Myr^-1^, an origination rate 0.8 Myr^-1^ and complete sampling in the present, i.e., *t* = 0 (ρ = 1) (Supplementary Table 3). Based on the extracted information on gaps (see above), we simulated three types of synthetic fossil record along the trees using the StratPal and FossilSim packages (Barido-Sottani, Pett, et al. 2019; Hohmann and Jarochowska 2025) with differing structures:

1. Constant fossil sampling rate (CFS), representing a fossil record that matches the assumptions of the FBD model.
2. Many short gaps (MSG) with no fossils preserved during gaps in the age-depth model taken from scenario 1 (Figure 1 C, Figure 2 A: Miller et al. sea level curve).
3. Few long gaps (FLG), with no fossils preserved during gaps in the age-depth model taken from scenario 2 (Figure 1 D, Figure 2:A Sinusoid sea level curve).

For this, we first simulated a large number of fossils under a constant sampling rate, removed those that coincided with gaps, and then uniformly subsampled the remaining ones to select 30 fossils to include in the inference. Fixing the number of preserved fossil isolates the effects of stratigraphic gaps and controls for different levels of information contributed by varying numbers of fossils. This setup focuses on structure (location and frequency of gaps) rather than (in)completeness (number of fossils recovered). Fossil sampling in the FBD model is determined by a Poisson point process, which is stable under random removal of events (thinning) (Streit 2010). Uniform subsampling will thus not introduce model violations. No fossil age uncertainty was incorporated (but see Barido-Sottani, Aguirre-Fernández, et al. (2019); Barido-Sottani et al. (2020)).

For extant taxa, a molecular partition of length 2000 was simulated under the HKY + G model with transition/transversion ratio of 5, 5 shape categories, and shape parameter of 0.25 following Barido-Sottani, Aguirre-Fernández, et al. (2019) using the phyclust package (Rambaut and Grass 1997; Chen and Dorman 2010) and assuming a strict clock model with clock rate 0.005 Myr^-1^. For all fossil and extant taxa, a morphological partition was simulated under the Mk(2) model (Lewis 2001) via the geiger package (Harmon et al. 2022) under a strict clock with rate 0.05 Myr^-1^ and 30, 100, 300, and 1000 characters, retaining noninformative characters.

### Phylogenetic inference

Total-evidence analysis was performed in RevBayes (Höhna et al. 2016) using three inference setups that represent increasing degrees of stratigraphic knowledge (Supplementary Table 4):

1. Stratigraphic agnosticism: No stratigraphic information is available and all rates are assumed to be constant throughout the examined time interval. Inference is run under the constant rate FBD (cFBD) model, with the inference setup matching the simulation setup under constant sampling. All parameters and the tree are estimated.
2. Weak stratigraphic knowledge: Inference is run under the “weak” skyline FBD (sFBDw) model, where knowledge of long gaps is incorporated by allowing sampling rates to vary between breakpoints taken from the stratigraphic forward model (scenario 2, sinusoidal sea level). All other parts of the inference match the simulation setup under constant sampling, all parameters (including per-interval fossil sampling rates) and the tree are estimated.
3. Strong stratigraphic knowledge: Inference is run under the “strong” skyline (sFBDs) model, where priors on fossil sampling are fixed to a very small numeric value during long gaps taken from the stratigraphic forward model. This formalizes the knowledge that no fossils can be sampled during these intervals. All other parts of the inference match the simulation setup under constant sampling, all parameters (except fossil sampling during gaps) and the tree are estimated.

Long gaps occurred only under the sinusoidal sea level (scenario 2) and were from 0.809 to 2.372 and 3.394 to 4.575 Ma at the sampling location (see above).

We ran inference under the constant rate FBD model on synthetic fossil records with constant fossil sampling (cFBD + CFS), many short gaps (cFBD + MSG) and few long gaps (cFBD + FLG) to examine how stratigraphically plausible model violations on fossil sampling affect phylogenetic inference. To explore the role of knowledge of gaps on phylogenetic inference, we ran inference under stratigraphic agnosticism (cFBD + FLG), weak stratigraphic knowledge (sFBDw + FLG), and strong stratigraphic knowledge (sFBDs + FLG) on a fossil record with few long gaps (FLG). Though run only once, the cFBD + FLG inference appears twice in our setup as it represents both stratigraphic agnosticism and inference under model violations.

For morphological and molecular clock rates and origination and extinction rates, we use exponential priors with the mean fixed to the true parameter value. For the origin time, we used an exponential prior with an offset of 5 Ma (true value) and a mean of 1 Myr. Exponential priors with means of 8 (cFBD), 5 (sFBDs) and 4 (sFBDw) were used for fossil sampling rate(s). See Supplementary Table 4 for a detailed description of priors. Due to the conditioning on the number of preserved fossils, the fossil sampling rates in cFBD and fossil sampling rates in sFBDw and sFBDs are considered nuisance parameters.

Inference was run for 100000 generations, with an additional burn-in interval of 20000 generations and parameter tuning for 15000 generations. Each generation consisted of 1570 (cFBD), 1630 (sFBDw) and 1600 (sFBDs) moves for the different degrees of gap knowledge. Two independent chains were run per analysis, with every 10^th^ generation saved. Runs were considered converged if all continuous parameters in both the merged and the individual runs had an effective sample size (ESS) above 200 (Supplementary Figure 3) (Barido-Sottani et al. 2024). Non-converged runs were not included in further analyses. A total of 40 replicates were run per inference setup, synthetic fossil record structure, and morphological character matrix size, resulting in 800 inferences.

### Summary statistics

For converged runs, we determined the relative error

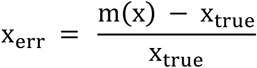

of model parameters use in the simulation setup (origination and extinction rate, molecular and morphological clock rate and time of origin). Here, *m*(*x*)is the median of the posterior sample of the merged trace of the two independent chains, and *x_true_* is the true value used for simulation (Figure 3, Figure 5). In addition, we determined the 90% credible interval coverage (frequency with which the true parameter is contained within the 90% credible interval) (Supplementary Figure 4) and the relative credible interval width (width of the 90 % credible interval divided by the true value) (Supplementary Figures 5 to 8). From merged tree traces, we determined the MAP tree and compared it to the simulated sampled tree (the “true” tree). We calculated (1) the RF distance (Robinson and Foulds 1981), (2) mean precision of divergence times between extant taxa (range of the 95% HDP interval divided by true divergence time) given the tree has two or more extant taxa, (3) 95% HDP coverage frequency of divergence times of extant taxa (proportion of divergences where the 95% HDP interval contains the true divergence time) given the tree has three or more extant taxa, and (4) the 95% sampled ancestor (SA) identification frequency (frequency with which fossils are correctly identified as SA or not). We considered a fossil correctly identified if its support for being a SA in the MAP tree is either (1) larger than 0.95 and it is a SA, or (2) below 0.05 and it is not an SA (Figure 4, Figure 6).

## Acknowledgements

We would like to thank Raoul Schram and Christine Staiger for their help with setting up workflows and data storage. We are indebted to Fiona and Shrek for their computational support.

## Code and data availability

All code is available under https://doi.org/10.5281/zenodo.20826148 (Hohmann and Jarochowska 2026), simulation and inference data is archived under https://doi.org/10.24416/UU01-6ON0ǪY (Hohmann 2026). Supplementary information is available under https://doi.org/10.5281/zenodo.20826528 (Hohmann, Warnock, et al. 2026)

## Author contributions

NH: Conceptualization, Formal analysis, Investigation, Methodology, Software, Validation, Visualization, Writing (original draft, review and editing). RW: Conceptualization, Writing (review and editing). EJ: Conceptualization, Funding acquisition, Validation, Supervision, Writing (review and editing)

## References

1. Alroy, John. 2002. “Stratigraphy in Phylogeny Reconstruction—Reply to Smith (2000).” Journal of Paleontology 76 (4): 587–89. 10.1666/0022-3360(2002)076%3C0587:SIPRRT%3E2.0.CO;2.

2. Barido-Sottani, J, O. Schwery, RCM Warnock, C. Zhang, and AM Wright. 2024. “Practical Guidelines for Bayesian Phylogenetic Inference Using Markov Chain Monte Carlo (MCMC) [Version 3; Peer Review: 4 Approved, 3 Approved with Reservations].” Open Research Europe 3 (204). 10.12688/openreseurope.16679.3.

3. Barido-Sottani, Joëlle, Gabriel Aguirre-Fernández, Melanie J. Hopkins, Tanja Stadler, and Rachel Warnock. 2019. “Ignoring Stratigraphic Age Uncertainty Leads to Erroneous Estimates of Species Divergence Times under the Fossilized Birth–Death Process.” Proceedings of the Royal Society B: Biological Sciences 286 (1902): 20190685. 10.1098/rspb.2019.0685.

4. Barido-Sottani, Joëlle, and Hélène Morlon. 2026. “Integrating Fossils Samples with Heterogeneous Diversification Rates: A Combined Multi-Type Fossilized Birth-Death Model.” *Systematic Biology*, February 17, syag011. 10.1093/sysbio/syag011.

5. Barido-Sottani, Joëlle, Walker Pett, Joseph E. O’Reilly, and Rachel C. M. Warnock. 2019. “FossilSim: An r Package for Simulating Fossil Occurrence Data under Mechanistic Models of Preservation and Recovery.” Methods in Ecology and Evolution 10 (6): 835–40. 10.1111/2041-210X.13170.

6. Barido-Sottani, Joëlle, Orlando Schwery, Rachel C. M. Warnock, Chi Zhang, and April Marie Wright. 2024. Using Markov Chain Monte Carlo (MCMC).

7. Barido-Sottani, Joëlle, Nina M. A. van Tiel, Melanie J. Hopkins, David F. Wright, Tanja Stadler, and Rachel C. M. Warnock. 2020. “Ignoring Fossil Age Uncertainty Leads to Inaccurate Topology and Divergence Time Estimates in Time Calibrated Tree Inference.” Frontiers in Ecology and Evolution 8 (June). 10.3389/fevo.2020.00183.

8. Benton, Michael J., and Brent C. Emerson. 2007. “How Did Life Become so Diverse? The Dynamics of Diversification According to the Fossil Record and Molecular Phylogenetics.” Palaeontology 50 (1): 23–40. 10.1111/j.1475-4983.2006.00612.x.

9. Bezanson, Jeff, Alan Edelman, Stefan Karpinski, and Viral B. Shah. 2017. “Julia: A Fresh Approach to Numerical Computing.” SIAM Review 59 (1): 65–98. 10.1137/141000671.

10. Capobianco, Alessio. 2025. “How Many Characters Are Needed to Reconstruct a Phylogeny?” Biology Letters 21 (10): 20250288. 10.1098/rsbl.2025.0288.

11. Catuneanu, Octavian. 2022. Principles of Sequence Stratigraphy. Newnes.

12. Chen, Wei-Chen, and Karin Dorman. 2010. “Phyclust: Phylogenetic Clustering (Phyloclustering).” January 29. 10.32614/CRAN.package.phyclust.

13. Danise, Silvia, and Steven M. Holland. 2017. “Faunal Response to Sea-Level and Climate Change in a Short-Lived Seaway: Jurassic of the Western Interior, USA.” Palaeontology 60 (2): 213–32. 10.1111/pala.12278.

14. Darwin, Charles. 1859. On the Origin of Species: A Facsimile of the First Edition. Harvard University Press.

15. Davies, Neil S., Anthony P. Shillito, and William J. McMahon. 2019. “Where Does the Time Go? Assessing the Chronostratigraphic Fidelity of Sedimentary Geological Outcrops in the Pliocene–Pleistocene Red Crag Formation, Eastern England.” Journal of the Geological Society 176 (6): 1154–68. 10.1144/jgs2019-056.

16. Didier, Gilles, Manuela Royer-Carenzi, and Michel Laurin. 2012. “The Reconstructed Evolutionary Process with the Fossil Record.” Journal of Theoretical Biology 315 (December): 26–37. 10.1016/j.jtbi.2012.08.046.

17. Donoghue, Philip C. J. 2001. “Conodonts Meet Cladistics: Recovering Relationships and Assessing the Completeness of the Conodont Fossil Record.” Palaeontology 44 (1): 65–93. 10.1111/1475-4983.00170.

18. Dzik, Jerzy. 2005. “The Chronophyletic Approach: Stratophenetics Facing an Incomplete Fossil Record.” Special Papers in Palaeontology 73.

19. Escapa, Ignacio H., and Diego Pol. 2011. “Dealing with Incompleteness: New Advances for the Use of Fossils in Phylogenetic Analysis.” PALAIOS 26 (3): 121–24. 10.2110/palo.2011.S02.

20. Fisher, Daniel C. 2008. “Stratocladistics: Integrating Temporal Data and Character Data in Phylogenetic Inference.” Annual Review of Ecology, Evolution, and Systematics 39 (Volume 39, 2008): 365–85. 10.1146/annurev.ecolsys.38.091206.095752.

21. Fisher, Daniel C., Michael Foote, David L. Fox, and Lindsey R. Leighton. 2002. “Stratigraphy in Phylogeny Reconstruction—Comment on Smith (2000).” Journal of Paleontology 76 (4): 585–86. 10.1666/0022-3360(2002)076%3C0585:SIPRCO%3E2.0.CO;2.

22. Flannery Sutherland, Joseph T., Benjamin C. Moon, Thomas L. Stubbs, and Michael J. Benton. 2019. “Does Exceptional Preservation Distort Our View of Disparity in the Fossil Record?” Proceedings of the Royal Society B: Biological Sciences 286 (1897): 20190091. 10.1098/rspb.2019.0091.

23. Hannisdal, Bjarte. 2006. “Phenotypic Evolution in the Fossil Record: Numerical Experiments.” The Journal of Geology 114 (2): 133–53. 10.1086/499569.

24. Heath, Tracy A., John P. Huelsenbeck, and Tanja Stadler. 2014. “The Fossilized Birth–Death Process for Coherent Calibration of Divergence-Time Estimates.” Proceedings of the National Academy of Sciences 111 (29): E2957–66. 10.1073/pnas.1319091111.

25. Hidding, J., E. Jarochowska, N. Hohmann, X. Liu, P. Burgess, and H. Spreeuw. 2025. “CarboKitten.Jl – an Open Source Toolkit for Carbonate Stratigraphic Modeling.” EGUsphere 2025: 1–28. 10.5194/egusphere-2025-4561.

26. Hohmann, Niklas. 2021. “Incorporating Information on Varying Sedimentation Rates into Paleontological Analyses.” PALAIOS 36 (2): 53–67. 10.2110/palo.2020.038.

27. Hohmann, Niklas. 2026. “Phylogenetic Inference from an Incomplete Fossil Record: Simulation and Inference Data.” Datapackage. Utrecht University, May 21. 10.24416/UU01-6ON0QY.

28. Hohmann, Niklas, Sidney Bickerton, Anna Jansen, Xianyi Liu, and Emilia Jarochowska. 2026. “Stratigraphic Paleobiology of Carbonate Systems.” Preprint, bioRxiv, June 17. 10.64898/2026.06.12.732006.

29. Hohmann, Niklas, David De Vleeschouwer, Sietske Batenburg, and Emilia Jarochowska. 2025. “Nonparametric Estimation of Age–Depth Models from Sedimentological and Stratigraphic Information.” Geochronology 7 (3): 427–48. 10.5194/gchron-7-427-2025.

30. Hohmann, Niklas, and Emilia Jarochowska. 2025. “StratPal: An R Package for Creating Stratigraphic Paleobiology Modelling Pipelines.” Methods in Ecology and Evolution 16 (4): 678–86. 10.1111/2041-210X.14507.

31. Hohmann, Niklas, and Emilia Jarochowska. 2026. Phylogenetic Inference from an Incomplete Fossil Record: Supplementary Code. V. v1.0.1. Zenodo, released June 24. 10.5281/ZENODO.20826148.

32. Hohmann, Niklas, Joël R. Koelewijn, Peter Burgess, and Emilia Jarochowska. 2024. “Identification of the Mode of Evolution in Incomplete Carbonate Successions.” BMC Ecology and Evolution 24 (1): 113. 10.1186/s12862-024-02287-2.

33. Hohmann, Niklas, Rachel Warnock, and Emilia Jarochowska. 2026. Phylogenetic Inference from an Incomplete Fossil Record: Supplementary Information. Version v1.0.0. June 24. 10.5281/ZENODO.20826528.

34. Höhna, Sebastian, Michael J. Landis, Tracy A. Heath, et al. 2016. “RevBayes: Bayesian Phylogenetic Inference Using Graphical Models and an Interactive Model-Specification Language.” Systematic Biology 65 (4): 726–36. 10.1093/sysbio/syw021.

35. Holland, Steven M. 2000. “The Quality of the Fossil Record: A Sequence Stratigraphic Perspective.” Paleobiology 26 (S4): 148–68. 10.1017/S0094837300026919.

36. Holland, Steven M. 2016. “The Non-Uniformity of Fossil Preservation.” Philosophical Transactions of the Royal Society B: Biological Sciences 371 (1699): 20150130. 10.1098/rstb.2015.0130.

37. Holland, Steven M. 2017. “Structure, Not Bias.” Journal of Paleontology 91 (6): 1315–17. 10.1017/jpa.2017.114.

38. Holland, Steven M., and Mark E. Patzkowsky. 1999. “Models for Simulating the Fossil Record.” Geology 27 (6): 491–94. 10.1130/0091-7613(1999)027%3C0491:MFSTFR%3E2.3.CO;2.

39. Holland, Steven M., and Mark E. Patzkowsky. 2015. “The Stratigraphy of Mass Extinction.” Palaeontology 58 (5): 903–24. 10.1111/pala.12188.

40. Holland, Steven M., Mark E. Patzkowsky, and Katharine M. Loughney. 2025. “Stratigraphic Paleobiology.” Paleobiology 51 (1): 44–61. 10.1017/pab.2024.2.

41. Kabanov, Pavel. 2017. “Stratigraphic Unconformities: Review of the Concept and Examples from the Middle-Upper Paleozoic.” In Seismic and Sequence Stratigraphy and Integrated Stratigraphy - New Insights and Contributions. IntechOpen. 10.5772/intechopen.70373.

42. Khakurel, Basanta, Courtney Grigsby, Tyler D. Tran, Juned Zariwala, Sebastian Höhna, and April M. Wright. 2024. “The Fundamental Role of Character Coding in Bayesian Morphological Phylogenetics.” Systematic Biology 73 (5): 861–71. 10.1093/sysbio/syae033.

43. Kidwell, Susan M., and Steven M. Holland. 2002. “The Quality of the Fossil Record: Implications for Evolutionary Analyses.” Annual Review of Ecology and Systematics 33 (1): 561–88. 10.1146/annurev.ecolsys.33.030602.152151.

44. Lewis, Paul O. 2001. “A Likelihood Approach to Estimating Phylogeny from Discrete Morphological Character Data.” Systematic Biology 50 (6): 913–25. 10.1080/106351501753462876.

45. Liu, Xianyi, Sam Purkis, Peter Burgess, et al. 2026. Stratigraphy as a Low-Pass Filter: Selective Preservation of Spatial Variability on a Holocene Carbonate Platform. May 20. https://eartharxiv.org/repository/view/13113/.

46. Loughney, Katharine M., and Steven M. Holland. 2026. “Simulations of Nonmarine Extensional Basins and Their Fossil Records.” Palaios 41 (2): 91–108. 10.2110/palo.2025.027.

47. Luke Harmon, Matthew Pennell, Chad Brock, Joseph Brown, Wendell Challenger, Jon Eastman, Rich FitzJohn, Rich Glor, Gene Hunt, Liam Revell, Graham Slater, Josef Uyeda, Jason Weir and CRAN team (corrections in 2022). 2007. “Geiger: Analysis of Evolutionary Diversification.” April 26. 10.32614/CRAN.package.geiger.

48. Magee, Andrew F., and Sebastian Höhna. 2021. “Impact of K-Pg Mass Extinction Event on Crocodylomorpha Inferred from Phylogeny of Extinct and Extant Taxa.” Preprint, bioRxiv, January 16. 10.1101/2021.01.14.426715.

49. Miller, Kenneth G., James V. Browning, W. John Schmelz, Robert E. Kopp, Gregory S. Mountain, and James D. Wright. 2020. “Cenozoic Sea-Level and Cryospheric Evolution from Deep-Sea Geochemical and Continental Margin Records.” Science Advances 6 (20): eaaz1346. 10.1126/sciadv.aaz1346.

50. Mulvey, Laura P. A., Mark C. Nikolic, Bethany J. Allen, Tracy A. Heath, and Rachel C. M. Warnock. 2025. “From Fossils to Phylogenies: Exploring the Integration of Paleontological Data into Bayesian Phylogenetic Inference.” Paleobiology 51 (1): 214–36. 10.1017/pab.2024.47.

51. Nawrot, Rafał, Daniele Scarponi, Michele Azzarone, et al. 2018. “Stratigraphic Signatures of Mass Extinctions: Ecological and Sedimentary Determinants.” Proceedings of the Royal Society B: Biological Sciences 285 (1886): 20181191. 10.1098/rspb.2018.1191.

52. Paul, Christopher R. C. 1992. “How Complete Does the Fossil Record Have to Be?” *Revista Española de Paleontología* 7 (2): 127–33.

53. R Core Team. 2025. R: A Language and Environment for Statistical Computing. R Foundation for Statistical Computing. https://www.R-project.org/.

54. Rambaut, Andrew, and Nicholas C. Grass. 1997. “Seq-Gen: An Application for the Monte Carlo Simulation of DNA Sequence Evolution along Phylogenetic Trees.” Bioinformatics 13 (3): 235–38. 10.1093/bioinformatics/13.3.235.

55. Robinson, D. F., and L. R. Foulds. 1981. “Comparison of Phylogenetic Trees.” Mathematical Biosciences 53 (1): 131–47. 10.1016/0025-5564(81)90043-2.

56. Ronquist, Fredrik, Seraina Klopfstein, Lars Vilhelmsen, Susanne Schulmeister, Debra L. Murray, and Alexandr P. Rasnitsyn. 2012. “A Total-Evidence Approach to Dating with Fossils, Applied to the Early Radiation of the Hymenoptera.” Systematic Biology 61 (6): 973–99. 10.1093/sysbio/sys058.

57. Sansom, Robert S., Emma Randle, and Philip C. J. Donoghue. 2015. “Discriminating Signal from Noise in the Fossil Record of Early Vertebrates Reveals Cryptic Evolutionary History.” Proceedings of the Royal Society B: Biological Sciences 282 (1800): 20142245. 10.1098/rspb.2014.2245.

58. Smith, Andrew B. 2000. “Stratigraphy in Phylogeny Reconstruction.” Journal of Paleontology 74 (5): 763–66. 10.1666/0022-3360(2000)074%3C0763:SIPR%3E2.0.CO;2.

59. Springer, Mark S. 1995. “Molecular Clocks and the Incompleteness of the Fossil Record.” Journal of Molecular Evolution 41 (5): 531–38. 10.1007/BF00175810.

60. Stadler, Tanja. 2010. “Sampling-through-Time in Birth–Death Trees.” Journal of Theoretical Biology 267 (3): 396–404. 10.1016/j.jtbi.2010.09.010.

61. Stadler, Tanja, Alexandra Gavryushkina, Rachel C. M. Warnock, Alexei J. Drummond, and Tracy A. Heath. 2018. “The Fossilized Birth-Death Model for the Analysis of Stratigraphic Range Data under Different Speciation Concepts.” arXiv:1706.10106. Preprint, arXiv, March 9. 10.48550/arXiv.1706.10106.

62. Straub, Kyle M., Robert A. Duller, Brady Z. Foreman, and Elizabeth A. Hajek. 2020. “Buffered, Incomplete, and Shredded: The Challenges of Reading an Imperfect Stratigraphic Record.” Journal of Geophysical Research: Earth Surface 125 (3): e2019JF005079. 10.1029/2019JF005079.

63. Streit, Roy L. 2010. Poisson Point Processes: Imaging, Tracking, and Sensing. Springer US. 10.1007/978-1-4419-6923-1.

64. Tanja Stadler. 2010. “TreeSim: Simulating Phylogenetic Trees.” February 23. 10.32614/CRAN.package.TreeSim.

65. Tipper, John C. 2015. “The Importance of Doing Nothing: Stasis in Sedimentation Systems and Its Stratigraphic Effects.” In Strata and Time: Probing the Gaps in Our Understanding, edited by D. G. Smith, R. J. Bailey, P. M. Burgess, and A. J. Fraser, vol. 404. Geological Society of London. 10.1144/SP404.6.

66. Tomašových, Adam, Ivo Gallmetzer, Alexandra Haselmair, and Martin Zuschin. 2022. “Inferring Time Averaging and Hiatus Durations in the Stratigraphic Record of High-Frequency Depositional Sequences.” Sedimentology 69 (3): 1083–118. 10.1111/sed.12936.

67. Turner, Alan H., Adam C. Pritchard, and Nicholas J. Matzke. 2017. “Empirical and Bayesian Approaches to Fossil-Only Divergence Times: A Study across Three Reptile Clades.” PLOS ONE 12 (2): e0169885. 10.1371/journal.pone.0169885.

68. Wagner, Peter J. 1995. “Stratigraphic Tests of Cladistic Hypotheses.” Paleobiology 21 (2): 153–78. 10.1017/S009483730001318X.

69. Wagner, Peter J., and Jonathan D. Marcot. 2013. “Modelling Distributions of Fossil Sampling Rates over Time, Space and Taxa: Assessment and Implications for Macroevolutionary Studies.” Methods in Ecology and Evolution 4 (8): 703–13. 10.1111/2041-210X.12088.

70. Williams, Huw D., Peter M. Burgess, V. Paul Wright, Giovanna Della Porta, and Didier Granjeon. 2011. “Investigating Carbonate Platform Types: Multiple Controls and a Continuum of Geometries.” Journal of Sedimentary Research 81 (1): 18–37. 10.2110/jsr.2011.6.

71. Woolley C. Henrik, David J. Bottjer, and Nathan D. Smith. 2025. “Taphonomic Megabiases Constrain Phylogenetic Information in the Squamate Fossil Record.” Paleobiology 51 (3): 554–73. 10.1017/pab.2025.10060.

72. Wright, April M., David W. Bapst, Joëlle Barido-Sottani, and Rachel C. M. Warnock. 2022. “Integrating Fossil Observations Into Phylogenetics Using the Fossilized Birth–Death Model.” *Annual Review of Ecology*, Evolution, and Systematics 53 (Volume 53, 2022): 251–73. 10.1146/annurev-ecolsys-102220-030855.

73. Zhang, Chi, Fredrik Ronquist, and Tanja Stadler. 2023. “Skyline Fossilized Birth–Death Model Is Robust to Violations of Sampling Assumptions in Total-Evidence Dating.” Systematic Biology 72 (6): 1316–36. 10.1093/sysbio/syad054.

74. Zhang, Chi, Tanja Stadler, Seraina Klopfstein, Tracy A. Heath, and Fredrik Ronquist. 2016. “Total-Evidence Dating under the Fossilized Birth–Death Process.” Systematic Biology 65 (2): 228–49. 10.1093/sysbio/syv080.

75. Zimmt, Joshua B., Steven M. Holland, Seth Finnegan, and Charles R. Marshall. 2021. “Recognizing Pulses of Extinction from Clusters of Last Occurrences.” Palaeontology 64 (1): 1–20. 10.1111/pala.12505.

